# A Next Generation Bivalent Human Ad5 COVID-19 Vaccine Delivering Both Spike and Nucleocapsid Antigens Elicits Th1 Dominant CD4+, CD8+ T-cell and Neutralizing Antibody Responses

**DOI:** 10.1101/2020.07.29.227595

**Authors:** Adrian Rice, Mohit Verma, Annie Shin, Lise Zakin, Peter Sieling, Shiho Tanaka, Helty Adisetiyo, Justin Taft, Rosheel Patel, Sofija Buta, Marta Martin-Fernandez, Brett Morimoto, Elizabeth Gabitzsch, Jeffrey T. Safrit, Joseph Balint, Kyle Dinkins, Patricia Spilman, Dusan Bogunovic, Shahrooz Rabizadeh, Kayvan Niazi, Patrick Soon-Shiong

## Abstract

In response to the health crisis presented by the COVID-19 pandemic, rapid development of safe and effective vaccines that elicit durable immune responses is imperative. Recent reports have raised the concern that antibodies in COVID-19 convalescent patients may not be long lasting and thus even these individuals may require vaccination. Vaccine candidates currently in clinical testing have focused on the SARS-CoV-2 wild type spike (S) protein (S-WT) as the major antigen of choice and while pre-clinical and early clinical testing have shown that S elicits an antibody response, we believe the optimal vaccine candidate should be capable of inducing robust, durable T-cell responses as well as humoral responses. We report here on a next generation bivalent human adenovirus serotype 5 (hAd5) vaccine capable of inducing immunity in patients with pre-existing adenovirus immunity, comprising *both* an S sequence optimized for cell surface expression (S-Fusion) and a conserved nucleocapsid (N) antigen designed to be transported to the endosomal subcellular compartment, with the potential to generate durable immune protection. Our studies suggest that this bivalent vaccine is optimized for immunogenicity as evidenced by the following findings: (i) The optimized S-Fusion displayed improved S receptor binding domain (RBD) cell surface expression compared to S-WT where little surface expression was detected; (ii) the expressed RBD from S-Fusion retained conformational integrity and recognition by ACE2-Fc; (iii) the viral N protein modified with an enhanced T-cell stimulation domain (ETSD) localized to endosomal/lysosomal subcellular compartments for MHC I/II presentation; and (iv) these optimizations to S and N (S-Fusion and N-ETSD) generated enhanced *de novo* antigen-specific B cell and CD4+ and CD8+ T-cell responses in antigen-naive pre-clinical models. Both the T-cell and antibody immune responses to S and N demonstrated a T-helper 1 (Th1) bias. The antibody responses were neutralizing as demonstrated by two independent SARS-CoV-2 neutralization assays. Based on these findings, we are advancing this next generation bivalent hAd5 S-Fusion + N-ETSD vaccine as our lead clinical candidate to test for its ability to provide robust, durable cell-mediated and humoral immunity against SARS-CoV-2 infection. Further studies are ongoing to explore utilizing this vaccine construct in oral, intranasal, and sublingual formulations to induce mucosal immunity in addition to cell-mediated and humoral immunity. The ultimate goal of an ideal COVID-19 vaccine is to generate long-term T and B cell memory.

## INTRODUCTION

In December of 2019, reports emerged from Wuhan, China concerning a new infectious respiratory disease with high morbidity and mortality^1-3^ that displayed human-to-human transmission.^4^ The causative agent was rapidly identified as a novel coronavirus and was designated SARS-coronavirus 2 (SARS-CoV-2). The disease it causes is referred to as COVID-19 and has rapidly become a worldwide pandemic that has disrupted socioeconomic life and resulted in more than 13 million infections and more than 600,000 deaths worldwide as of late July 2020.^5,6^

SARS-CoV-2 is an enveloped positive sense, single-strand RNA β coronavirus primarily composed of four structural proteins - spike (S), nucleocapsid (N), membrane (M), and envelope (E) – as well as the viral membrane and genomic RNA. Of these, S is the largest and N the most prevalent. The S glycoprotein is displayed as a trimer on the viral surface (Fig. 1a), whereas N is located within the viral particle. A schematic of the S primary structure is shown in Fig. 1b.^7^ The sequence of SARS-CoV-2 was published^8^ and compared to that of previous coronaviruses.^9,10^ This was soon followed by reports on the crystal structure of the S protein.^7,11^ The virus uses S protein to enter host cells by interaction of the S receptor binding domain (S RBD) with angiotensin-converting enzyme 2 (ACE2), an enzyme expressed broadly on a variety of cell types in the nose, mouth, gut and lungs as well as other organs, and importantly on the alveolar epithelial cells of the lung where infection is predominantly manifested.^12^ As represented in Fig 1b, the S RBD is found within the S1 region of spike.^7^

**Fig. 1.**
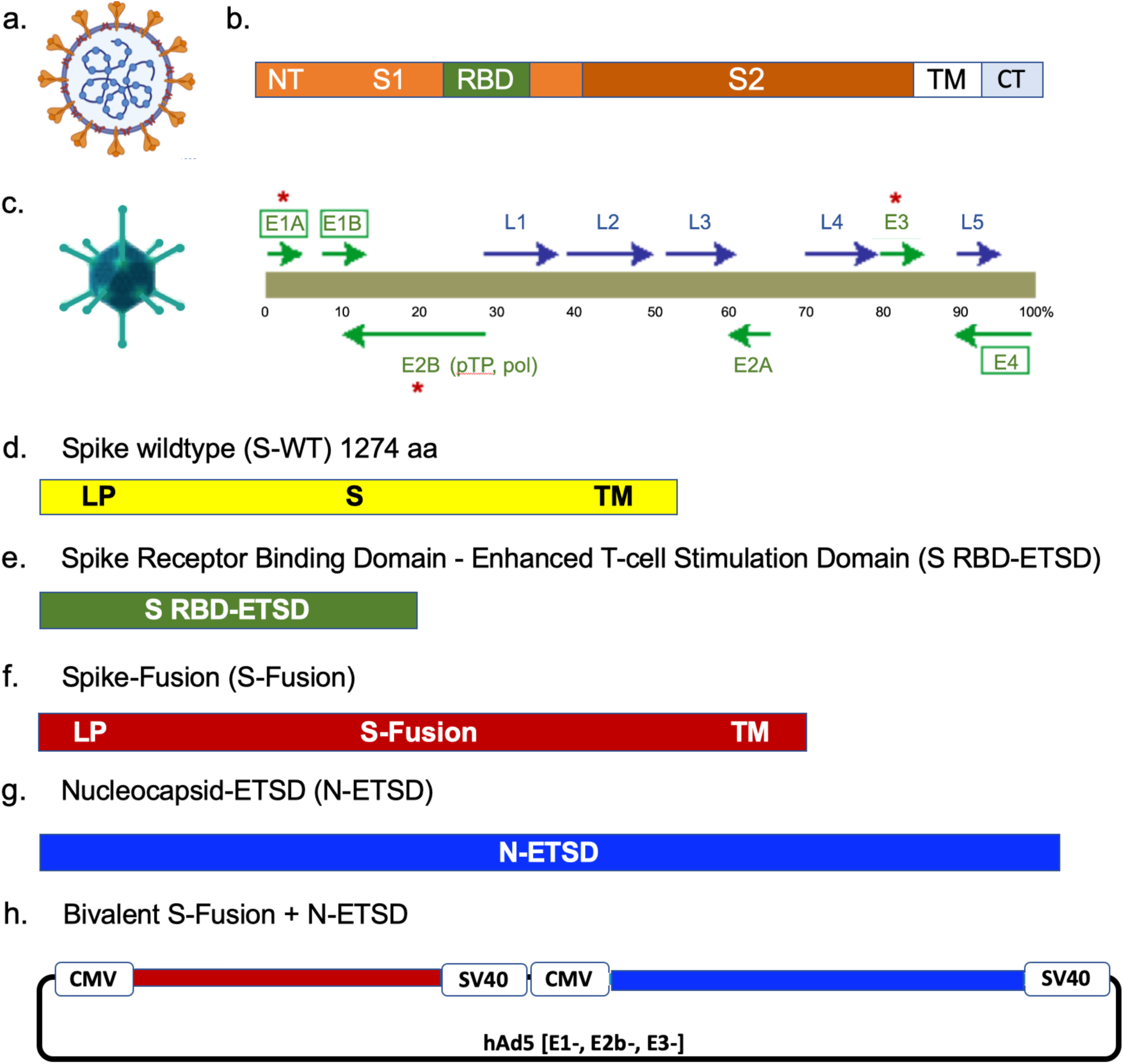
The SARS-CoV-2 virus, spike, the hAd5 [E1-, E2b-, E3-] vector and vaccine candidate constructs. (a) Trimeric spike (S) protein (▾) is displayed on the viral surface; the nucleocapsid (N) protein (○) is associated with the viral RNA. (b) The Receptor Binding Domain (RBD) is within the S1 region, followed by other functional regions, the transmembrane domain (TM) and the C-terminus (CT), which is within the virus. (c) The second-generation human adenovirus serotype 5 (hAd5) vector used has the E1, E2b, and E3 regions deleted. Constructs are shown for (d) S wild type (S-WT), (e) S-RBD with the Enhanced T-cell Stimulation Domain (S RBD-ETSD), (f) S-Fusion, (g) N-ETSD, and (h) bivalent hAd5 S-Fusion + N-ETSD; LP – Leader peptide.

The majority of current SARS-CoV-2 vaccines under development target S because of the potential to neutralize the ability of the virus to bind host cells by production of antibodies against the RBD.^13^ Support for RBD as a key antigen was recently confirmed by Suthar *et al*.,^14^ who reported that in 44 hospitalized COVID-19 patients, RBD-specific IgG responses and neutralizing antibody titers are detectable in all patients by 6 days post-PCR confirmation of infection, and that the two are correlated. They confirmed this finding in an additional 231 PCR-confirmed COVID-19 patient samples. In addition to humoral responses, S epitopes are also frequent targets of COVID-19 recovered patient T cells,^15^ providing further justification for inclusion of S in prophylactic immunization strategies.

Despite the urgent need for rapid development of SARS-CoV-2 vaccines, reliance on any one antigen cargo or immunological pathway as occurring in the monovalent vaccines under development is not without risk. Evaluation of nearly 4000 SARS-CoV-2 genomic sequences has identified numerous mutations in S (https://nextstrain.org/ncov/global) with the D614G variant emerging recently as a potentially more infectious strain six months after identification of the original virus.^16^

In our vaccine design, to overcome the risk of the emergence of new strains of the virus with mutations in S and to provide additional antigens against which responses can be elicited, we have added an optimized N sequence. The N protein is a highly conserved and antigenic SARS-CoV-2-associated protein that has been studied previously as an antigen in coronavirus vaccine design for SARS-CoV.^17-20^ N associates with viral RNA within the virus and has a role in viral RNA replication, virus particle assembly, and release.^21,22^ SARS-CoV-2 N is a highly antigenic protein and recent studies have shown that nearly all patients infected with SARS-CoV-2 have antibody responses to N.^23,24^ Furthermore, another study reported that most, if not all, COVID-19 survivors tested were shown to have N-specific CD4+ T-cell responses.^15^

Currently, there is keen focus on generation of humoral responses to vaccines with, arguably, less attention being paid to T-cell responses. The natural history of SARS-CoV-2 infection would suggest, however, that a robust T-cell response to vaccination is at least as important as the production of antibodies ^25^ and should be a critical consideration for COVID-19 vaccine efficacy. First, the humoral and T-cell responses are highly correlated, with titers of neutralizing antibodies being proportional to T-cell levels, suggesting the T response is necessary for an effective humoral response.^26^ It is well established that the activation of CD4+ T helper cells enhances B-cell production of antibodies. Second, virus-specific CD4+ and CD8+ T cells are not only widely detected in COVID-19 patients,^27^ based on findings from patients recovered from the closely-related SARS-CoV, but such T cells persist for at least 6–17 years, suggesting that T cells may be an important part of long-term immunity.^28-30^ These T-cell responses were predominantly to N, as described in Le Bert *et al*., who found that in all 36 convalescent COVID-19 patients in their study, the presence of CD4+ and CD8+ T cells recognizing multiple regions of the N protein could be demonstrated.^30^ They further examined blood from 23 individuals who had recovered from SARS-CoV and found that the memory T cells acquired 17 years ago also recognized multiple proteins of SARS-CoV-2. These findings emphasize the importance of designing a vaccine with the highly conserved nucleocapsid present in both SARS-CoV and SARS-CoV-2. Third, recovered patients exposed to SARS-CoV-2 have been found without seroconversion, but with evidence of T-cell responses.^31^ The T-cell based responses become even more critical given the finding in at least one study that neutralizing antibody titers decline in some COVID-19 patients after about 3 months.^32^

Thus, we have paid particular attention in the design of our vaccine to the generation of T-cell in addition to humoral responses. It is our hypothesis that a bivalent vaccine comprising many antigens – S RBD as displayed by inclusion of full-length S including SD1, S1 and S2 epitopes, along with N – could be more effective in eliciting both T-cell and antibody-based responses than a construct with either antigen alone by presenting both unique and conserved SARS-CoV-2 antigenic sites to the immune system. The importance of both S and N was highlighted by Grifoni *et al*.^15^ who identified both S and N antigens as *a priori* potential B and T-cell epitopes for the SARS-CoV virus that shows close similarity to SARS-CoV-2 that are predicted to induce both T and B cell responses.

An additional consideration for design of an effective vaccine is the likelihood of antigen presentation on the surface of the vectored-protein-expressing cell and in a conformation that recapitulates natural virus infection. First, because wild type N does not have a signaling domain that directs it to endosomal processing and ultimately MHC class II complex presentation to CD4+ T cells, the wild type N sequence is not optimal for induction of a vigorous CD4+ T-cell responses, a necessity for both cell-mediated and B cell memory. To overcome this limitation, we have designed an Enhanced T-cell Stimulation Domain (ETSD) to N to allow the necessary processing and presentation. Second, to display the highly antigenic RBD region of S on the cell surface, we have optimized the wild type S protein “S Fusion sequence”, to increase the likelihood of native folding, increased stability, and proper cell surface expression of RBD. Our innovative vaccine construct design comprises an S-Fusion + N-ETSD sequence.

The vaccine platform utilized here is a next-generation recombinant human adenovirus serotype 5 (hAd5) vector with deletions in the E1, E2b, and E3 gene regions (hAd5 [E1-, E2b-, E3-]).^33^ This hAd5 [E1-, E2b-, E3-] vector (Fig. 1c) is primarily distinguished from other first-generation [E1-, E3-] recombinant Ad5 platforms ^34,35^ by having additional deletions in the early gene 2b (E2b) region that remove the expression of the viral DNA polymerase (pol) and in pre terminal protein (pTP) genes, and its propagation in the E.C7 human cell line.^33,36-38^ Removal of these E2b regions confers advantageous immune properties by minimizing immune responses to Ad5 viral proteins such as viral fibers, ^37^ thereby eliciting potent immune responses to specific antigens in patients with pre-existing adenovirus (Ad) immunity. ^39,40^ As a further benefit of these deletions, the vector has an expanded gene-carrying/cloning capacity compared to the first generation Ad5 [E1-, E3-] vectors. This next generation hAd5 [E1-, E2b-, E3-] vaccine platform, in contrast to Ad5 [E1-, E3-]-based platforms, does not promote activities that suppress innate immune signaling, thereby allowing for improved vaccine efficacy and a superior safety profile independent of previous Ad immunity.^41^ Since these deletions allow the hAd5 platform to be efficacious even in the presence of existing Ad immunity, this platform enables relatively long-term antigen expression without significant induction of anti-vector immunity. It is therefore also possible to use the same vector/construct for homologous prime-boost therapeutic regimens unlike first-generation Ad platforms which face the limitations of pre-existing and vaccine-induced Ad immunity.^42^ Importantly, this next generation Ad vector has demonstrated safety in over 125 patients with solid tumors. In these Phase I/II studies, CD4+ and CD8+ antigen-specific T cells were successfully generated to multiple somatic antigens (CEA, MUC1, brachyury) even in the presence of pre-existing Ad immunity.^39 43^

Herein, we report our findings of confirmed enhanced cell-surface expression and physiologically-relevant folding of the expressed S RBD from S-Fusion by ACE2-Fc binding. We report that the N-ETSD protein successfully localized to the endosomal/lysosomal subcellular compartment for MHC presentation and consequently generated both CD4+ and CD8+ T-cell responses. Immunization of CD-1 mice with the hAd5 S Fusion + N-ETSD vaccine elicited both humoral and cell-mediated immune responses to vaccine antigens. CD8+ and CD4+ T-cell responses were noted for both S and N. Statistically significant IgG responses were seen for antibody generation against S and N. Potent neutralization of SARS-CoV-2 by sera from hAd5 S Fusion + N-ETSD-immunized mice was confirmed by two independent SARS-CoV-2 neutralization assays: the cPass assay measuring competitive inhibition of RBD binding to ACE2,^44^ and in the live SARS-CoV-2 virus assay with infected Vero E6 cells. Analysis of T-cell responses as well as humoral responses to S and N were skewed toward a Th1-specific response. Taken together, these findings have led to the decision that hAd5 S-Fusion + N-ETSD is our lead candidate to advance to clinical trials.

## RESULTS

### The hAd5 [E1-, E2b-, E3-] platform and constructs

For studies here, the next generation hAd5 [E1-, E2b-, E3-] vector was used (Fig. 1c) to create viral vaccine candidate constructs. As shown in Figure 1d-h, a variety of constructs were created:

d. S WT: S protein comprising 1273 amino acids and all S domains: extracellular (1-1213), transmembrane (1214-1234), and cytoplasmic (1235-1273) (Unitprot P0DTC2);

e. S RBD-ETSD: S Receptor Binding Domain with an Enhanced T-cell Stimulation Domain (ETSD);

f. S Fusion: S optimized to enhance surface expression and display of RBD;

g. N-ETSD: The nucleocapsid (N) sequence with the ETSD; and

h. Bivalent S-Fusion + N-ETSD;

S-WT + N-ETSD and S RBD-ETSD + N-ETSD constructs were also produced, but are not shown.

### Enhanced HEK 293T cell-surface expression of RBD following transfection with Ad5 S-Fusion + N-ETSD

As shown in Figure 2, anti-RBD-specific antibodies did not detect RBD on the surface of HEK 293T cells transfected with hAd5 S-WT (Fig. 2a) or hAd5 S-WT + N-ETSD (Fig. 2b) constructs, while hAd5 S-Fusion alone was slightly higher (Fig. 2e). As expected, both constructs with RBD, hAd5 RBD-ETSD and RBD-ETSD + N-ETSD, showed high binding of anti-RBD antibody (Fig. 2c and d). Notably, high cell-surface expression of RBD was detected after transfection with bivalent hAd5 S-Fusion + N-ETSD (Fig. 2f). These findings support our proposition that an hAd5 S-Fusion + N-ETSD construct, containing a high number and variety of antigens provided by both full-length, optimized S with proper folding and N leads to enhanced expression and cell surface display of RBD in a vaccine construct.

**Fig. 2.**
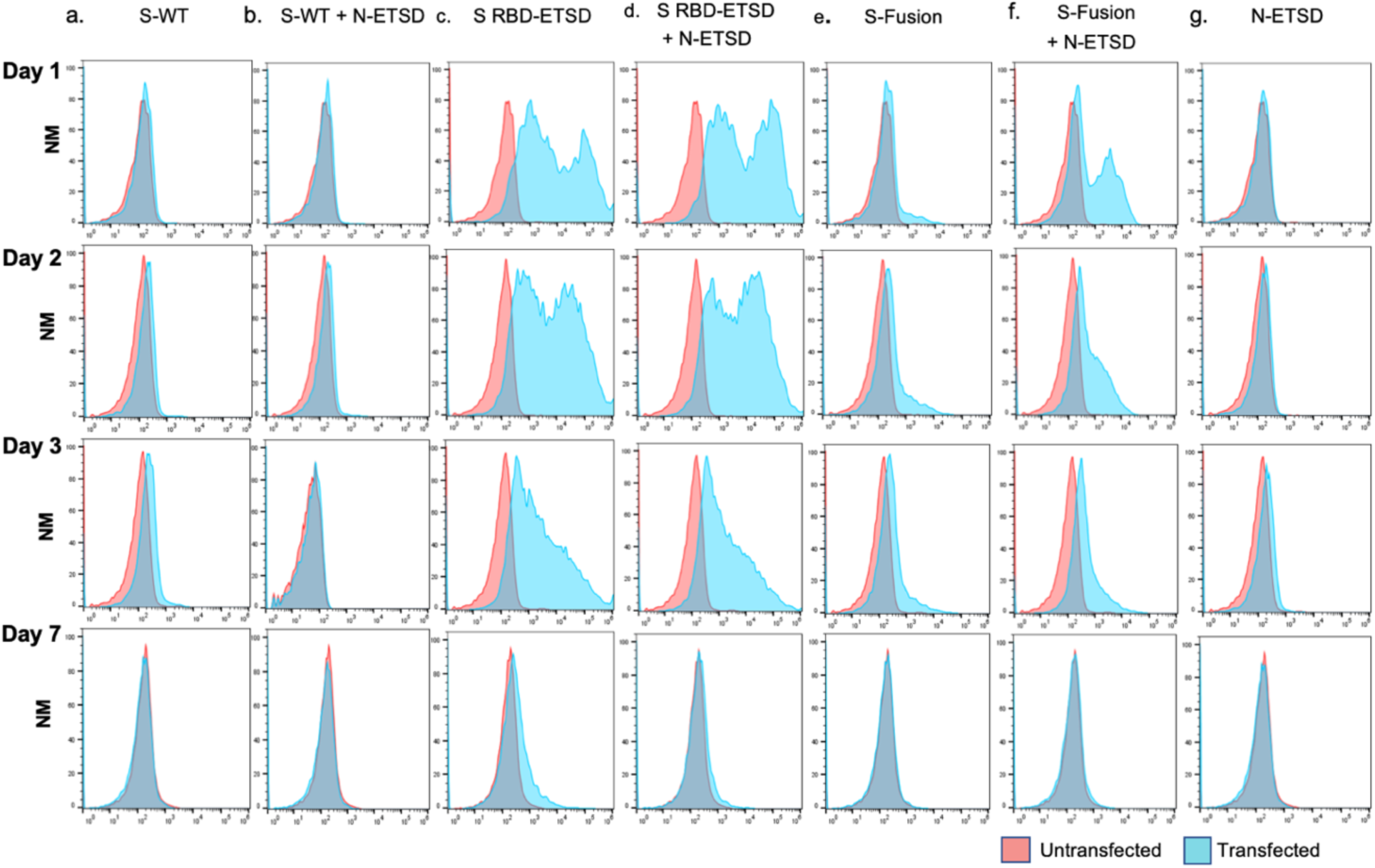
Transfection of HEK293T cells with hAd5 S-Fusion + ETSD results in enhanced surface expression of the spike receptor binding domain (RBD). Flow cytometric analysis of an anti-RBD antibody with construct-transfected cells reveals no detectable surface expression of RBD in either (a) S-WT or (b) S-WT + N-ETSD transfected cells. Surface RBD expression was high for S RBD-ETSD and S RBD-ETSD + N-ETSD (c, d). Expression was low in (e) S-Fusion transfected cells. Cell surface expression of the RBD was high in (f) S-Fusion + N-ETSD transfected cells, particularly at day 1 and 2. (g) No expression was detected the N-ETSD negative control. Y-axis scale is normalized to mode (NM).

### Immunoblot correlation of enhanced S expression with hAd5 S-Fusion + N-ETSD

Immunoblot analysis of S expression correlated with enhanced S expression (Fig. 3), showing again that the bivalent hAd5 S-Fusion + N-ETSD construct enhances expression of S compared to S-Fusion alone.

**Fig. 3.**
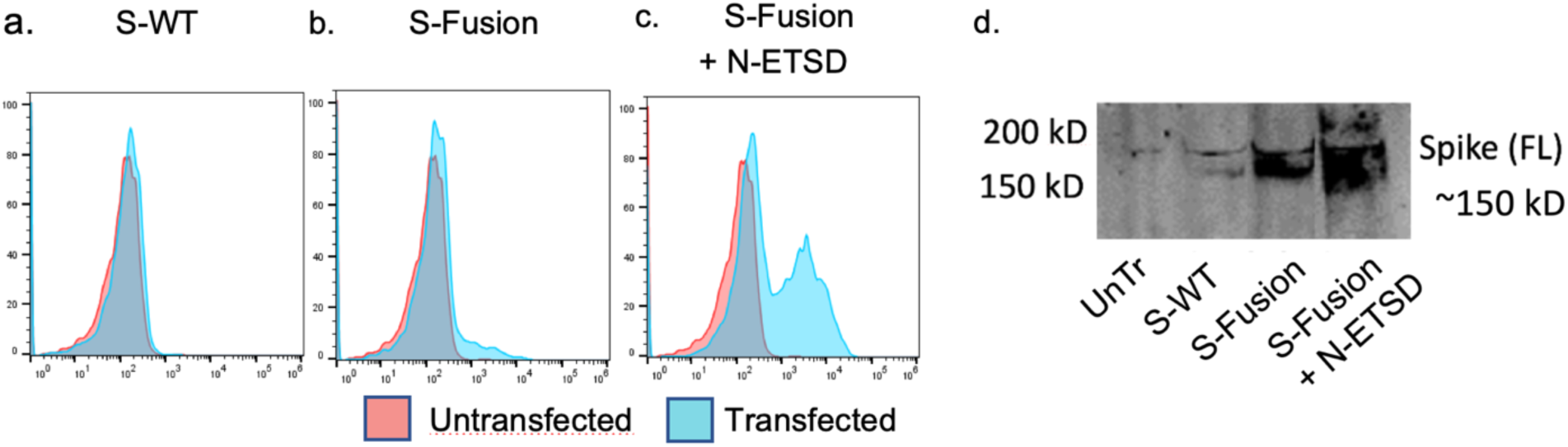
Immunoblot analysis of S expression. Cell surface RBD expression with (a) hAd5 S-WT, (b) S-Fusion, and (c) S-Fusion + N-ETSD in HEK 293T cells shows high correlation with (d) expression of S in immunoblots of HEK 293T cell lysates probed using anti-full length (S2) antibody. Y-axis scale is normalized to mode (NM).

### Confirmation of native folding of enhanced surface RBD following hAd5 S-Fusion + N-ETSD transfection

Determination of the binding of recombinant ACE2-Fc was performed to confirm the native, physiologically-relevant folding of the S RBD after expression from the hAd5 S-Fusion +N-ETSD vaccine candidate. S RBD binds ACE2 during the course of SARS-CoV-2 infection and an effective neutralizing antibody prevents this interaction and thus infection. Such a neutralizing antibody is more likely to be effective if raised in response to S presented in the correct conformation. In addition to enhancement of cell surface expression, the optimized S allows for proper protein folding. We found that compared to either hAd5 S-WT or hAd5 S-Fusion (Fig. 4 and b, respectively), ACE2-Fc binding to S RBD expressed from the hAd5 S-Fusion + N-ETSD was clearly enhanced (Fig. 4c). Anti-RBD antibody binding studies (Fig. 4f-j) performed with the same experiment, confirmed the enhanced surface expression findings noted by ACE2-Fc binding. These findings of conformationally correct and enhanced S RBD expression, important for production of neutralizing antibodies, directed us to elect the hAd5 S-Fusion + N-ETSD vaccine candidate for clinical trials.

**Fig. 4.**
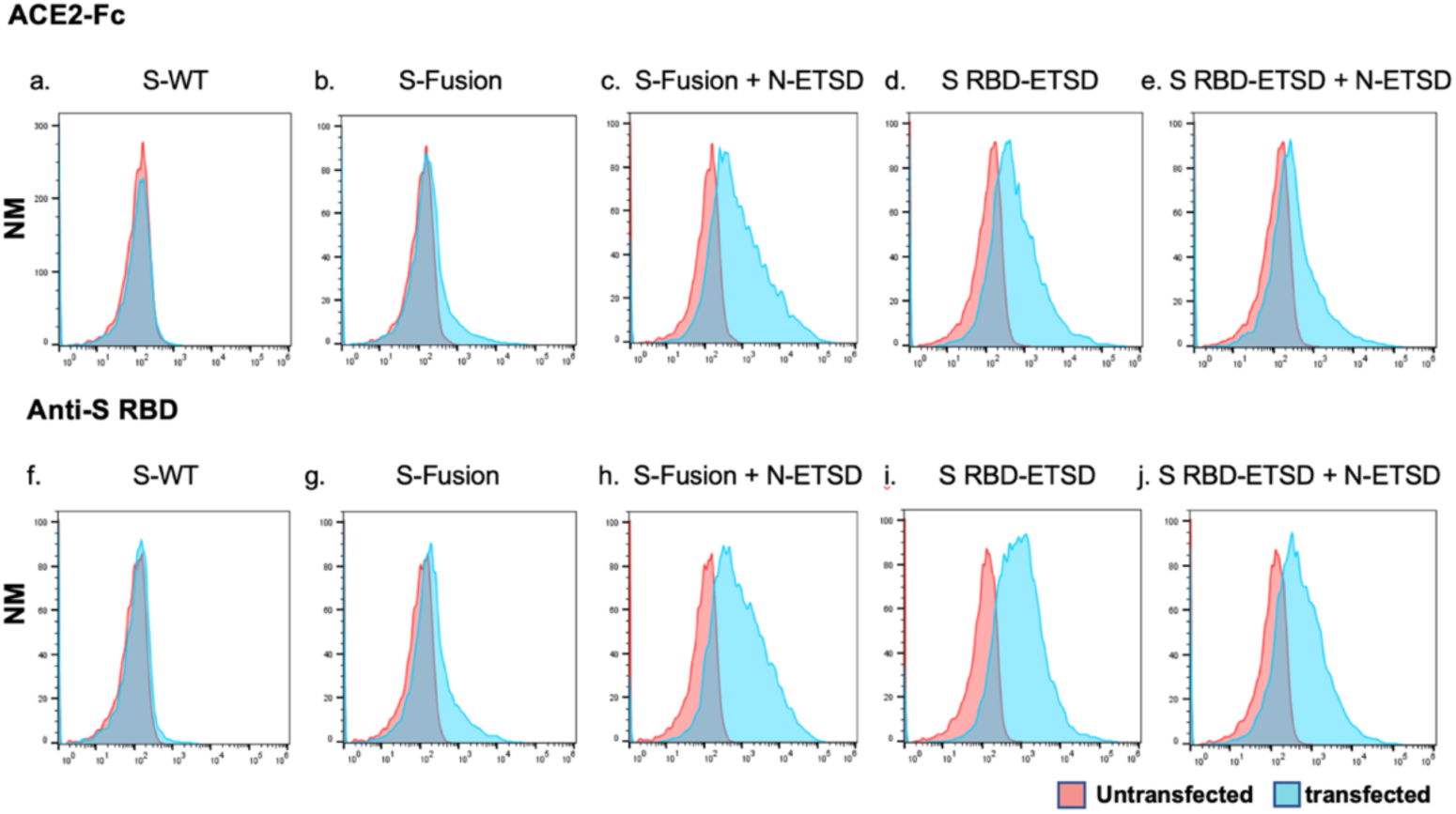
Binding of recombinant ACE2-Fc HEK293T cell-surface expressed RBD after transfection confirms native protein folding. Flow cytometric analysis of binding between recombinant ACE2-Fc, with which the spike RBD interacts *in vivo* to initiate infection, and cell-surface antigens expressed after transfection of HEK293T cells with (a) hAd5 S-WT, (b) hAd5 S-Fusion, (c) hAd5 S-Fusion + N-ETSD, (d) hAd5 S RBD-ETSD, or (e) hAd5 S RBD-ETSD + N-ETSD constructs reveals the highest binding is seen for both ACE-Fc and an anti-RBD specific antibody (f-j) after transfection with the bivalent S-Fusion + N-ETSD. Both S RBD-ETSD-containing constructs also showed binding. Y-axis scale is normalized to mode (NM).

### hAd5 N-ETSD successfully directs N to an endosomal/lysosomal compartment

Our ETSD design successfully translocated N to the endosomal subcellular compartment. After infection of HeLa cells with N-ETSD, N co-localized with the endosomal marker ^45^ transferrin receptor (CD71), as shown in Fig. 5c, and also co-localized with the lysosomal marker Lamp1 (Fig. 5d), demonstrating that N-ETSD is translocated throughout the endosomal pathway to lysosomes, enabling processing for MHC II presentation. N-wild type (N-WT), compared to N-ETSD, shows diffuse cytoplasmic distribution and does not co-localize with the lysosomal marker (Fig. 5e). These findings confirm the role of the ETSD in directing N to an endosomal/lysosomal compartment that will result in increased MHC II presentation and CD4+ activation by N.

**Fig. 5.**
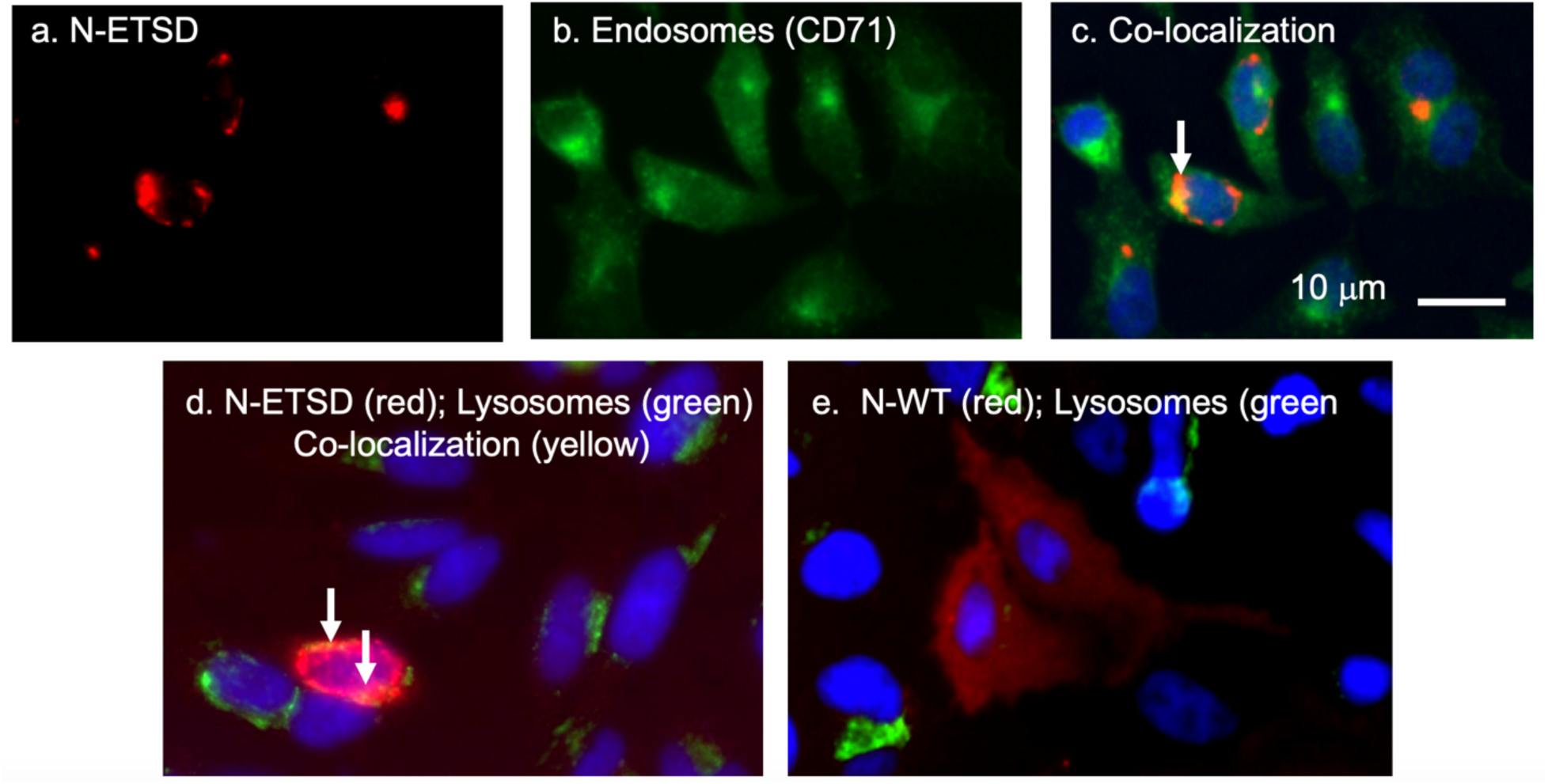
N expressed from hAd5 N-ETSD is localized to the endosomal/lysosomal compartment. In HeLa cells infected with N-ETSD, (a) N (red) co-localizes with the endosomal marker CD71 (b) as indicated by the arrow in (c). In transfected HeLa cells, (d) N-ETSD also co-localizes with the lysosomal marker Lamp1, whereas (e) N wild type (N-WT) does not, showing instead diffuse cytoplasmic distribution.

### *In Vivo* hAd5 S-Fusion + N-ETSD Vaccine Immunogenicity Studies

Based on the evidence that S-Fusion + N-ETSD resulted in enhanced expression of physiologically-relevant RBD and that N-ETSD successfully translocated to the endosomal / lysosomal compartment, we chose the bivalent hAd5 S-Fusion + N-ETSD vaccine for inoculation of 7-week old female CD-1 mice. We hypothesized that the unique properties of this construct would result in the generation of both CD8+ and CD4+ T-cell responses and neutralizing antibodies. As described in *Methods*, mice received an initial injection on Day 0 and a second injection on Day 21. Sera were collected on Day 0 and at the end of the study on Day 28 for antibody and neutralization analyses. Splenocytes were also collected on Day 28 for intracellular cytokine staining (ICS) and ELISpot analyses. All age- and gender-matched animals assigned to the study appeared normal with no site reactions and no loss of body weight throughout the dosing were seen, consistent with previous observations with the hAd5 [E1-, E2b-, E3-] platform.

### hAd5 S-Fusion + N-ETSD generates both CD8β+ and CD4+ T-cell responses

#### CD8+ activation by both S and N

CD8β^+^ splenocytes from hAd5 S-Fusion + N-ETSD vaccinated mice exposed to S peptide pool 1 (containing RBD and S1) show IFN-γ expression that is significantly higher compared to hAd5 null mice (Fig. 6a); splenocytes from these mice also expressed intracellular IFN-γ in response to the N peptide pool. Evaluation of simultaneous IFN-γ/TNF-α expression from CD8β+ splenocytes (Fig. 6c) mirrored those for IFN-γ expression alone. These results indicate that both S and N activate CD8+ T cells.

**Fig. 6.**
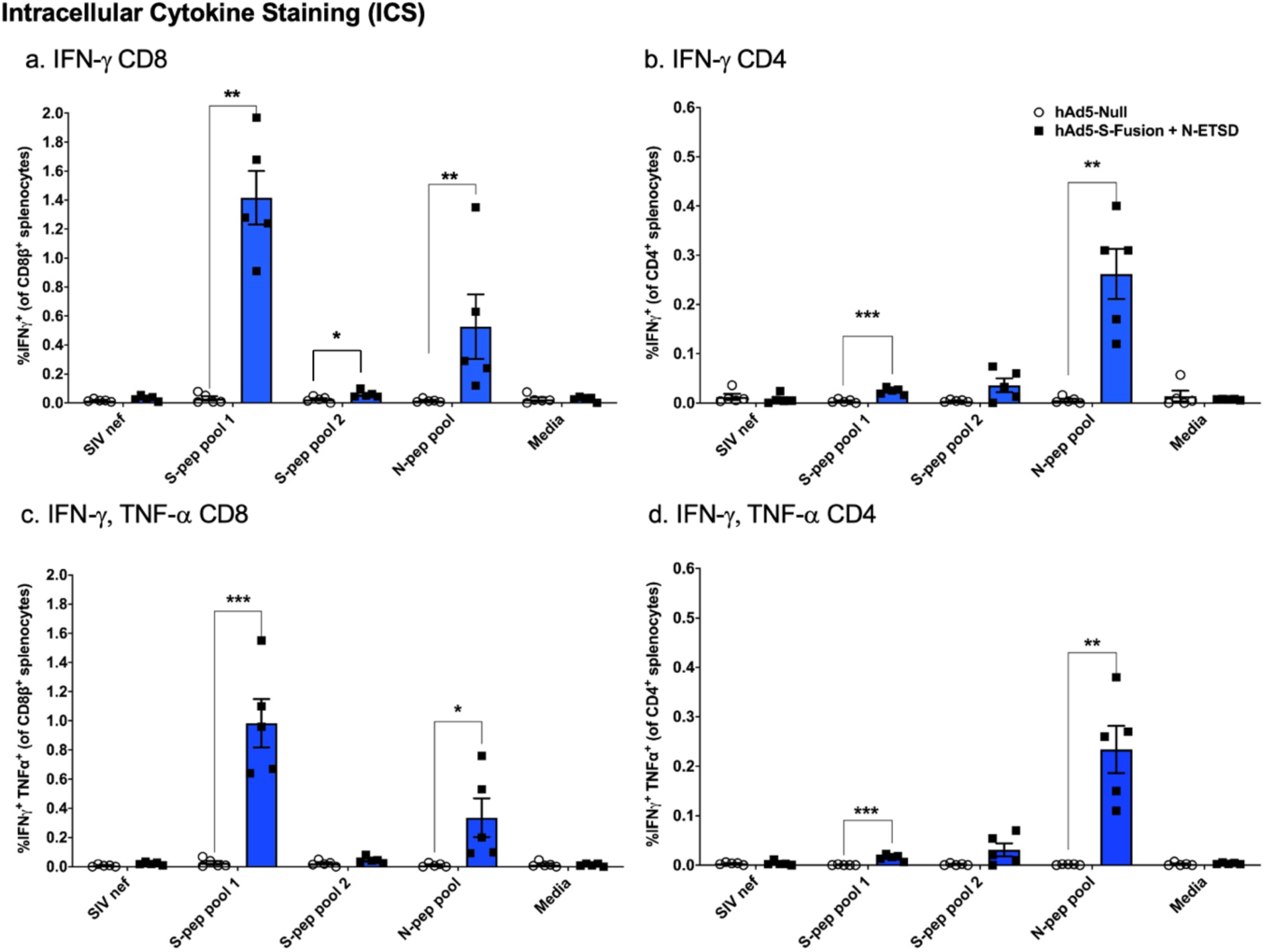
ICS detection of cytokine-expressing splenocytes from hAd5 S-Fusion + N-ETSD inoculated Day 28 CD-1 mice in response to peptide pools. (a) The highest CD8β^+^ splenocyte IFN-γ response was in hAd5 S-Fusion + N-ETSD-inoculated mouse splenocytes exposed to S peptide pool 1 (S-pep pool 1); splenocytes from these mice also expressed IFN-γ in response to the N peptide pool (N-pep pool). (b) CD4+ splenocytes from hAd5 S-Fusion + N-ETSD-inoculated mice only expressed IFN-γ in response to the N peptide pool. (c) IFN-γ TNF-α responses of CD8β^+^ splenocytes from hAd5 S-Fusion + N-ETSD-inoculated mice were very similar to those in (a); as were (d) CD4+ splenocytes to the N peptide pool to those in (b). N = 5 mice per group. All data sets graphed as the mean with SEM and all statistics performed using the Mann-Whitney test where *<0.05, **<0.01, ***<0.001, and ****<0.0001.

#### CD4+ activation by N

Although CD8+ cytotoxic T cells mediate killing of virus infected cells, CD4+ T cells are required for sustained cytotoxic T lymphocyte (CTL) activity.^46^ Thus we evaluated CD4+ T cells in the vaccinated animals. In contrast to CD8β+ splenocytes, only the N peptide pool stimulated CD4+ splenocytes from hAd5 S-Fusion + N-ETSD-inoculated mice to express IFN-γ (Fig. 6b) or IFN-γ/TNF-α (Fig. 6d) at levels that were substantially higher than hAd5 Null control. The contribution by N of CD4+ T-cell responses is vital to an effective immune response to the candidate vaccine.

### hAd5 S-Fusion + N-ETSD generates antibody responses to both S and N antigens

The primary objective of coronavirus vaccines currently in development are neutralizing antibodies against spike, thus we examined antibody production in mice vaccinated with our bivalent vaccine. There was significant production of both anti-S (Fig. 7a) and anti-N (Fig. 7c) antibodies in the sera from CD-1 mice vaccinated with hAd5 S-Fusion + N-ETSD at Day 28 in the study. Compared to anti-S antibodies, anti-N antibodies were higher in sera, given the dilution factor for sera was 1:90 for anti-N antibody analysis and 1:30 for anti-S antibody analysis.

**Fig. 7.**
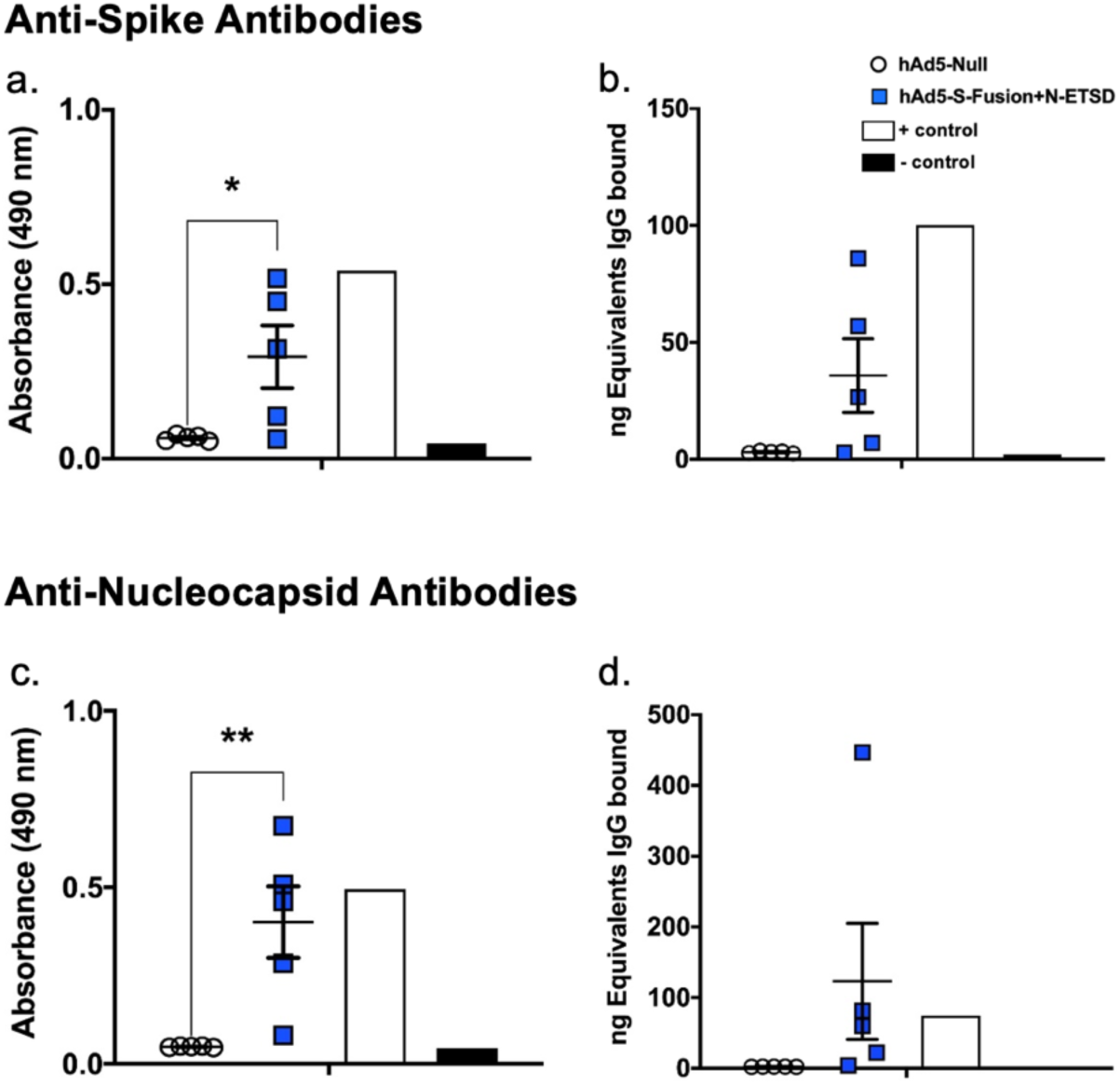
Anti-spike and anti-nucleocapsid antibody responses in sera from hAd5 S-Fusion + N-ETSD vaccinated mice. Based on absorbance, there was significant production of both (a) anti-S antibodies and (c) anti-nucleocapsid antibodies. (b, d) The ng equivalents are shown. Sera diluted 1:30 for anti-spike and 1:90 for anti-nucleocapsid antibodies. Data graphed as the mean and SEM. Statistical analysis was performed using an unpaired two-tailed Student’s t-test where *<0.05, **<0.01, ***<0.001, and ****<0.0001.

A standard curve of IgG was generated, then absorbance values were converted into mass equivalents for both anti-S and anti-N antibodies (Fig. 7b and d). These values were used to calculate that hAd5 S-Fusion + NETSD vaccination generated a geometric mean value of 5.8 μg S-specific IgG and 42 μg N-specific IgG per mL of serum, therefore the relative μg amount of anti-N antibodies is higher than that for anti-S antibodies and reflects the strong contribution of N to anti-SARS-CoV-2 antibody production.

### hAd5 S-Fusion + N-ETSD vaccine generates potent neutralizing antibodies as assessed by both cPass and live virus neutralization assays

Neutralizing antibody activity was evaluated using a cell free assay (cPass) as well as live virus infection *in vitro*. As seen in Figure 8a, the cPass assay showed inhibition of S RBD:ACE2 binding for all mice and ∼100% inhibition for two mice at both dilutions of 1:20 and 1:60. The Vero E6 neutralization assay results are shown for the four mice that showed S-specific antibodies by ELISA. The high persistent neutralization seen even at the high dilution factors suggests the intriguing possibility that the bivalent, multi-antigen, multi-epitope generation by hAd5 S-Fusion + N-ETSD vaccine, could result in synergies of neutralizing immune responses (Fig. 8b); at epitopes in addition to those associated with RBD-ACE2 binding. As can be seen in Fig. 8b, the value for 50% neutralization (IC50) is present at 1:10,000 serum dilution for the G4 pool of sera from mice that showed S-specific antibodies, ten times higher than the convalescent serum with a dilution of 1:1,000. The potent neutralization, confirmed by two assays, supports the predicted efficacy of the hAd5 S-Fusion + ETSD vaccine candidate and its advancement to clinical trials.

**Fig. 8.**
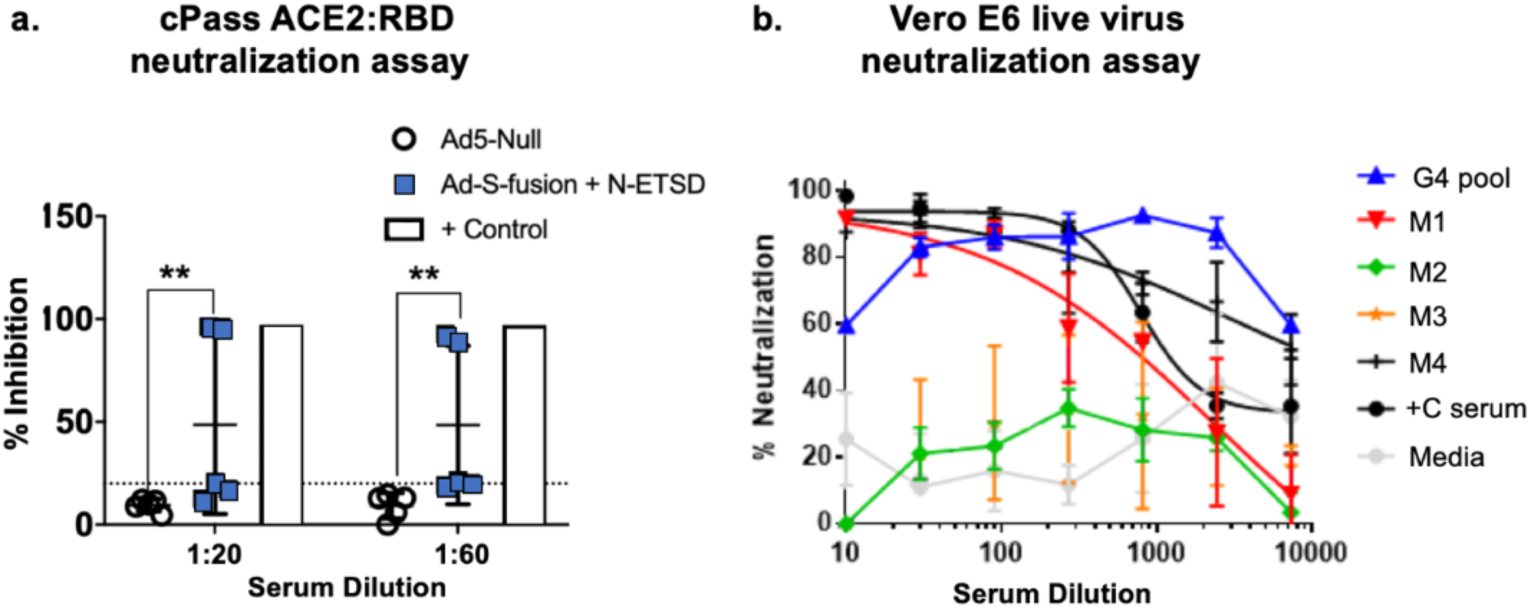
cPass and Vero E6 cell SARS-CoV-2 confirm neutralization by antibodies. (a) In the cPass assay, inhibition of S RBD interaction with ACE2 was significant at both 1:20 and 1:60 dilutions of serum from hAd5 S-Fusion + N-ETSD vaccinated mice. (b) The results in the Vero E6 cell SARS-CoV-2 viral infection for mice that showed S-specific antibodies by ELISA also showed high neutralization for mice and very high neutralization for pooled sera (G4 pool, blue line) even compared to COVID-19 convalescent serum. G4 pool – mice with S-specific antibodies; M1, M2, M3, M4 – mouse ID; +C – convalescent serum; and media – media only negative control.

### hAd5 S-Fusion + N-ETSD generates Th1 dominant responses both in humoral and T-cell immunity

#### Antibody Th1 dominance in response to N and S

IgG2a, IgG2b, and IgG3 represent Th1 dominance; while IgG1 represents Th2 dominance. For both anti-S (Fig, 9a) and anti-N (Fig, 9c) antibodies in sera from hAd5 S-Fusion + N-ETSD vaccinated mice, IgG2a and IgG2b isotypes were predominant and significantly higher compared to the hAd5 Null control. These data show the Th1 dominance of antibody production in response to the hAd5 S-Fusion + N-ETSD vaccine.

**Fig. 9.**
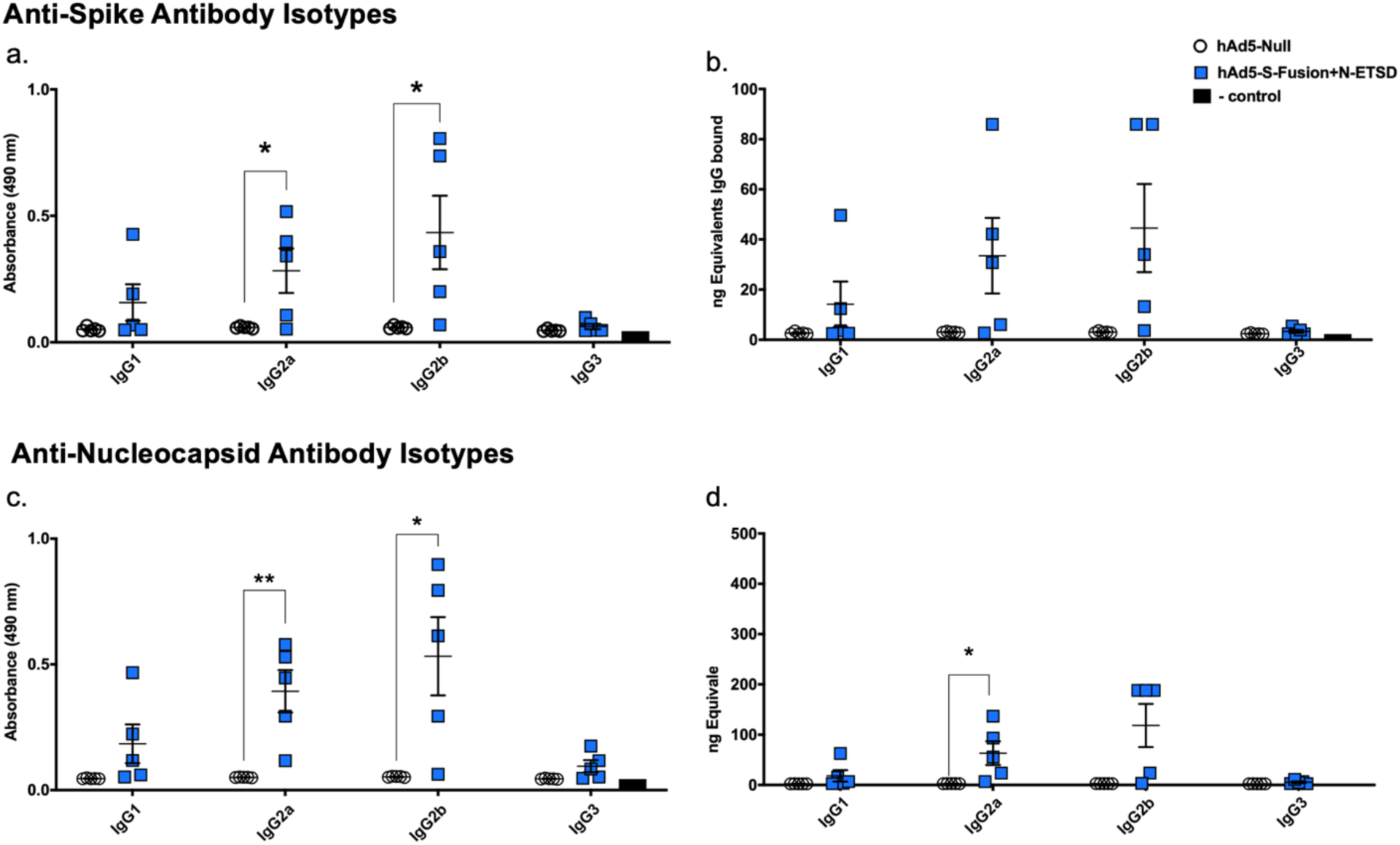
Isotypes for anti-spike and anti-nucleocapsid antibodies. (a, c) IgG2a and IgG2b isotype anti-spike and anti-nucleocapsid antibodies were significantly increased for hAd5 S-Fusion + N-ETSD mice compared to hAd5 Null mice. (b, d) The ng equivalents for antibody isotypes are shown. Data graphed as the mean and SEM. Statistical analysis was performed using an unpaired two-tailed Student’s t-test where *<0.05, **<0.01, ***<0.001, and ****<0.0001.

#### T-cell Th1 dominance in response to N and S

IFN-γ production correlates with CTL activity ^47^ (Th1 dominance), whereas, IL-4 causes delayed viral clearance ^48^ (Th2 dominance). A ratio of IFN-γ to IL-4 of 1 is balanced and a ratio greater than 1 is demonstrative of Th1 dominance. Thus, we examined IFN-γ and IL-4 production in animals immunized with the bivalent S plus N vaccine. As determined by ELISpot, IFN-γ secretion was significantly higher for hAd5 S-Fusion + N-ETSD than for hAd5 Null splenocytes in response to both S peptide pool 1 and the N peptide pool (Fig. 10a), but IL-4 was only secreted at significantly higher levels for hAd5 S-Fusion + N-ETSD in response to the N peptide pool (Fig. 10b).

**Fig. 10.**
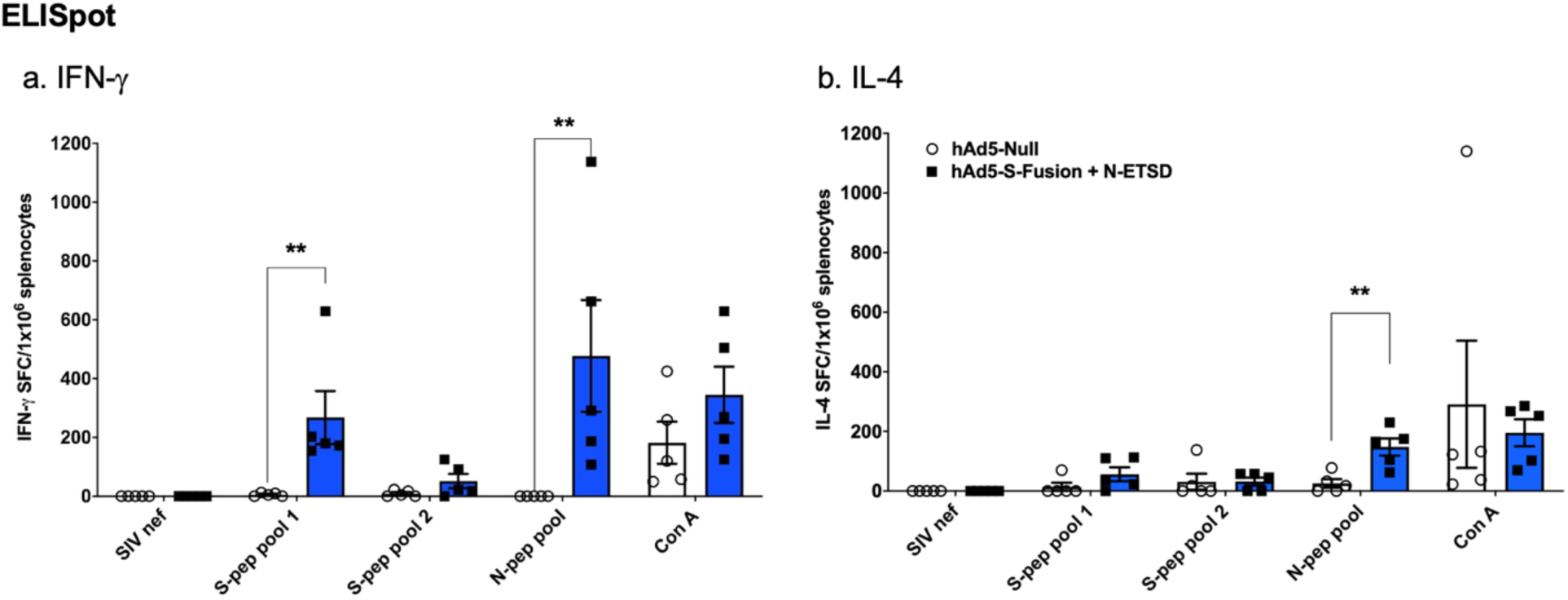
ELISpot detection of secreted cytokines. (a) IFN-γ secretion by hAd5 S-Fusion + N-ETSD splenocytes was significantly higher than hAd5 Null in response to both S peptide pool 1 and the N peptide pool; but (b) IL-4 was only secreted with hAd5 S-Fusion + N-ETSD in response to the N peptide pool (one high outlier in hAd5 null removed). N = 5 mice per group. All data sets graphed as the mean with SEM and all statistics performed using the Mann-Whitney test where *<0.05, **<0.01, ***<0.001, and ****<0.0001.

The Th1-type predominance is also seen when the ratio of IFN-γ to IL-4 based on spot forming units in response to the combined S peptide pools and the N peptide pool, is considered (Fig. 11a).

**Fig. 11.**
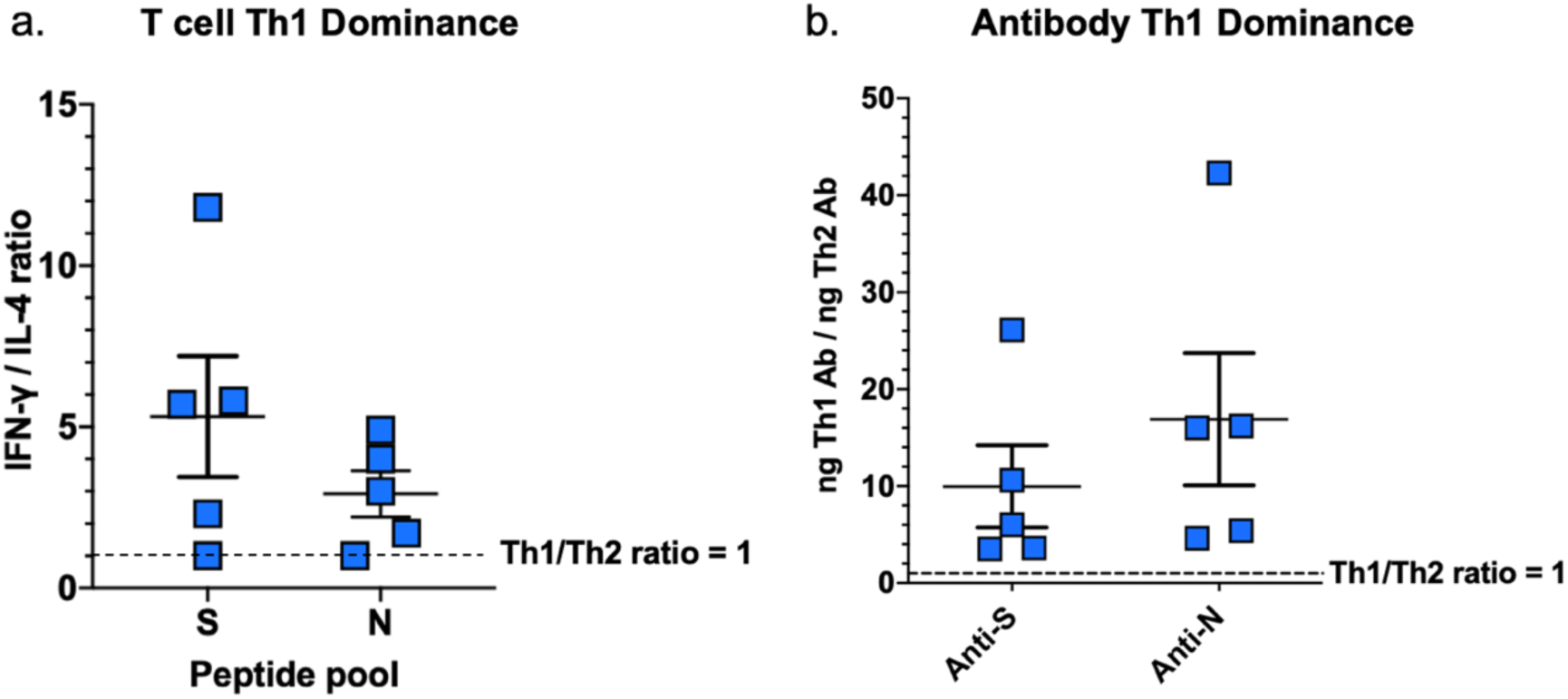
Ratios for T-cell and humoral responses reveal Th1 predominance. (a) The ratio of total Th1 (IFN-γ) to Th2 (IL-4) spot-forming units is shown for responses to the combined S pools and to the N pool. (b) The Th1/Th2 ratio for antibodies against S and N is shown. For both (a) and (b) the dashed line indicates a ratio of 1 or a balance of Th1 and Th2 (no predominance).

Th1 predominance was seen again in humoral responses, where the ratio based on ng equivalence of Th1 related antibodies (IgG2a, IgG2b, and IgG3) to Th2 related antibodies (IgG1) for both anti-S and anti-N antibodies is greater than 1 in all mice (Fig. 11b).

This Th1 dominant profile of the hAd5 S-Fusion + N-ETSD vaccine candidate provides further justification for hAd5 S-Fusion + N-ETSD to be our lead candidate for clinical testing.

## DISCUSSION

Our hAd5 S-Fusion + N-ETSD vaccine was designed to overcome the risks of an S-only vaccine and elicit both T-cell immunity and neutralizing antibodies, leveraging the vital role T cells play in generating long-lasting antibody responses and in directly killing infected cells. Both CD4+ and CD8+ T cells are multifunctional, and induction of such multifunctional T cells by vaccines correlated with better protection against infection.^49^ We posit that enhanced CD4+ T-cell responses and Th1 predominance resulting from expression of an S antigen optimized for surface display and an N antigen optimized for endosomal/lysosomal subcellular compartment localization and thus MHC I and II presentation, led to increased dendritic cell presentation, cross-presentation, B cell activation, and ultimately high neutralization capability. Furthermore, the potent neutralization capability at high dilution seen for the pooled sera from hAd5 S-Fusion + N-ETSD vaccinated mice, combined with Th1 dominance of antibodies generated in response to both S and N antigens, supports the objective of this vaccine design.

It is well established that the contemporaneous MHC I and MHC II presentation of an antigen by the antigen presenting cell activates CD4+ and CD8+ T cells simultaneously and is optimal for the generation of memory B and T cells. A key finding of our construct is that N-ETSD, which we show is directed to the endosomal/lysosomal compartment, elicits a CD4+ response, a necessity for induction of memory T cells and helper cells for B cell antibody production. Others have also reported on the importance of lysosomal localization for eliciting the strongest T-cell IFN-γ and CTL responses, compared to natural N.^50,51^

The T-cell responses to the S and N antigens expressed by hAd5 S-Fusion + N-ETSD were polycytokine, including IFN-γ and TNF-α, consistent with successful antimicrobial immunity in bacterial and viral infections.^52-56^ Post-vaccination polycytokine T-cell responses have been shown to correlate with vaccine efficacy, including those with a viral vector.^49^ Highly relevant here, polycytokine T-cell responses to SARS-CoV-2 N protein are consistent with recovered COVID-19 patients,^17^ suggesting that the bivalent hAd5 S-Fusion + N-ETSD vaccine will provide vaccine subjects with greater protection against SARS-CoV-2.

In contrast to N, the S protein, here expressed as S-Fusion with confirmed enhanced RBD cell-surface expression and conformational integrity as evidenced by high ACE2-Fc binding, generated predominantly CD8+ T cells. Our results confirmed our vaccine design goal, showing that S-Fusion induced elevated levels of antigen-specific T-cell responses against S compared to S-WT. To ensure MHC presentation to both MHC I (for CD8+ T-cell activation) and MHC II (for CD4+ T-cell activation), we believe it is necessary to vaccinate with both S and N antigens optimized to produce this coordinated response.

Our neutralization data with live SARS-CoV-2 virus demonstrated the potency of the antibody response generated following vaccination with hAd5 S-Fusion + N-ETSD, with evidence of high neutralization even at a high dilution factor. In addition, a striking synergistic effect of pooled sera was evident, with potent neutralization even greater than control convalescent serum at ≥ 1:1,000 dilution.

The hAd5 S-Fusion + N-ETSD construct described above is delivered by a next generation hAd5 [E1-, E2b-, E3-] platform wherein the E2b deletion (pol) alone enables prolonged transgene production^57^ and allows homologous vaccination (prime and the boost formulation is the same) in the presence of pre-existing adenoviral immunity.^38^ In addition to the generation of cellular and humoral immunity by the subcutaneous injection of hAd5 S-Fusion + N-ETSD, we are also exploring the potential of inducing IgA mucosal immunity by utilizing the same vaccine in an oral or sublingual formulation in clinical trials.

## METHODS

### The hAd5 [E1-, E2b-, E3-] platform and constructs

For studies herein, the 2^nd^ generation hAd5 [E1-, E2b-, E3-] vector was used (Fig. 1c) to create viral vaccine candidate constructs. hAd5 [E1-, E2b-, E3-] backbones containing SARS-CoV-2 antigen expressing inserts and virus particles were produced as previously described.^37^ In brief, high titer adenoviral stocks were generated by serial propagation in the E1- and E2b-expressing E.C7 packaging cell line, followed by CsCl2 purification, and dialysis into storage buffer (2.5% glycerol, 20 mM Tris pH 8, 25 mM NaCl) by ViraQuest Inc. (North Liberty, IA). Viral particle counts were determined by sodium dodecyl sulfate disruption and spectrophotometry at 260 and 280 nm and viral titers were determined using the Adeno-X™ Rapid Titer Kit (Takara Bio). The constructs created included:

i. S-WT: S protein comprising 1273 amino acids and all S domains: extracellular (1-1213), transmembrane (1214-1234), and cytoplasmic (1235-1273) (Unitprot P0DTC2);
ii. S RBD-ETSD: S Receptor Binding Domain (S RBD) with an ETSD;
iii. N-ETSD: Nucleocapsid (N) with ETSD;
iv. S-WT + N-ETSD: S-WT with an Enhanced T-cell Stimulation Domain (ETSD);
v. S-RBD-ETSD + N-ETSD;
vi. S Fusion: S optimized to enhance surface expression and display of RBD; and
vii. Bivalent S-Fusion + N-ETSD;

### Transfection of HEK 293T cells with hAd5 constructs

To determine surface expression of the RBD epitope by vaccine candidate constructs, we transfected HEK 293T cells with hAd5 construct DNA and quantified surface RBD by flow cytometric detection using anti-RBD antibodies. There were seven constructs tested: S-WT, S-WT + N-ETSD, S RBD-ETSD, S RBD-ETSD + N-ETSD, S-Fusion, S-Fusion + N-ETSD, and N-ETSD. HEK 293T cells (2.5 × 10^5^ cells/well in 24 well plates) were grown in DMEM (Gibco Cat# 11995-065) with 10% FBS and 1X PSA (100 units/mL penicillin, 100 μg/mL streptomycin, 0.25 ug/mL Amphotericin B) at 37°C. Cells were transfected with 0.5 μg of hAd5 plasmid DNA using a JetPrime transfection reagent (Polyplus Catalog # 89129-924) according to the manufacturer’s instructions. Cells were harvested 1, 2, 3, and 7 days post transfection by gently pipetting cells into medium and labeled with an anti-RBD monoclonal antibody (clone D003 Sino Biological Catalog # 40150-D003) and F(ab’)2-Goat anti-Human IgG-Fc secondary antibody conjugated with R-phycoerythrin (ThermoFisher Catalog # H10104). Labeled cells were acquired using a Thermo-Fisher Attune NxT flow cytometer and analyzed using Flowjo Software.

### Immunocytochemical labeling of hAd5 infected HeLa cells

To determine subcellular localization of N after infection or transfection of HeLa cells with hAd5 N-wild type (WT) or hAd5 N-ETSD (each with a flag tag to allow labeling), 48 hours after infection or transfection cells were fixed with 4% paraformaldehyde (PFA) and permeabilized with 0.4% Triton X100, in PBS) for 15 min. at room temperature. To label N, cells were then incubated with an anti-flag monoclonal (Anti-Flag M2 produced in mouse, Sigma cat# F1804) antibody at 1:1000 in phosphate buffered saline with 3% BSA overnight at 4°C, followed by washes in PBS and a 1 hour incubation with a goat anti-Mouse IgG (H+L) Highly Cross-Adsorbed Secondary Antibody, Alexa Fluor Plus 555 (Life Technologies, Cat# A32727) at 1:500. For co-localization studies, cells were also incubated overnight at 4°C with a sheep anti-Lamp1 Alexa Fluor 488-conjugated (lysosomal marker) antibody (R&D systems, Cat# IC7985G) at 1:10 or a rabbit anti-CD71 (transferrin receptor, endosomal marker) antibody (ThermoFisher Cat# PA5-83022) at 1:200. After removal of the primary antibody, two washes in PBS and three 3 washes in PBS with 3% BSA, cells were incubated with fluor-conjugated secondary antibodies when applicable at 1:500 (Goat anti-Rabbit IgG (H+L) Highly Cross-Adsorbed Secondary Antibody, Alexa Fluor 488, Life technologies, A-11034) for 1 hour at room temperature. After brief washing, cells were mounted with Vectashield Antifade mounting medium with DAPI (Fisher Scientific, Cat#NC9524612) and immediately imaged using a Keyence all-in-one Fluorescence microscope camera and Keyence software.

### Immunoblot analysis of S antigen expression

HEK 293T cells transfected with hAd5 S-WT, S-Fusion, or S-Fusion + N-ETSD constructs were cultured and transfected as described in the main manuscript and harvested 3 days after transfection in 150 mL RIPA lysis buffer with 1X final Protease Inhibitor cocktail (Roche). After protein assay, equivalent amounts of total protein were loaded into and run on a 4 to 12% gradient polyacrylamide gel (type) and transferred to nitrocellulose membranes using semi-dry transfer apparatus. Anti-Spike S2 (SinoBiological Cat #40590-T62) was used as the primary antibody and IRDye® 800CW Goat anti-Rabbit IgG (H + L) (Li-Cor, 925-32211) as the secondary antibody using the Ibind Flex platform. Antibody-specific signals were detected with an infrared Licor Odyssey instrument.

### ACE2-IgG1Fc binding to hAd5 transfected HEK 293T cells

HEK 293T cells were cultured at 37°C under conditions described above for transfection with hAd5 S-WT, S-Fusion, S-Fusion + N-ETSD, S RBD-ETSD, or S RBD-ETSD + N-ETSD and were incubated for 2 days and harvested for ACE2-Fc binding analysis. Recombinant ACE2-IgG1Fc protein was produced using Maxcyte transfection in CHO-S cells that were cultured for 14 days. ACE2-IgG1Fc was then purified using a MabSelect SuRe affinity column on AKTA Explorer. Purified ACE2-IgG1Fc was dialyzed into 10 mM HEPES, pH7.4, 150 mM NaCl and concentrated to 2.6 mg/mL. For binding studies, the ACE2-IgG1Fc was used at a concentration of 1 μg/mL for binding. Cells were incubated with ACE2-Fc for 20 minutes and, after a washing step, were then labeled with a PE conjugated F(ab’)2-goat anti-human IgG Fc secondary antibody at a 1:100 dilution, incubated for 20 minutes, washed and acquired on flow cytometer. Histograms are based on normalized mode (NM) of cell count – count of cells positive for signal in PE channel.

### Vaccination of CD-1 mice with the hAd5 S-Fusion + N-ETSD vaccine candidate

CD-1 female mice (Charles River Laboratories) 7 weeks of age were used for immunological studies performed at the vivarium facilities of Omeros Inc. (Seattle, WA). After an initial blood draw, mice were injected with either hAd5 Null (a negative control) or vaccine candidate hAd5 S-Fusion + N-ETSD on Day 0 at a dose of 1 ×10^10^ viral particles (VP). There were 5 mice per group. Mice received a second vaccine dose on Day 21 and on Day 28, blood was collected via the submandibular vein from isoflurane-anesthetized mice for isolation of sera and then mice were euthanized for collection of spleen and other tissues.

### Splenocyte collection and Intracellular cytokine staining (ICS)

Spleens were removed from each mouse and placed in 5 mL of sterile medium of RPMI (Gibco Cat # 22400105), HEPES (Hyclone Cat# SH30237.01), 1X Pen/Strep (Gibco Cat # 15140122), and 10% FBS (Gibco Cat # 16140-089). Splenocytes were isolated within 2 hours of collection. ICS for flow cytometric detection of CD8β^+^ and CD4^+^ T-cell-associated IFN-γ and IFN-γ/TNFα+ production in response to stimulation by S and N peptide pools.

Stimulation assays were performed using 10^6^ live splenocytes per well in 96-well U-bottom plates. Splenocytes in RPMI media supplemented with 10% FBS were stimulated by the addition of peptide pools at 2 μg/mL/peptide for 6 h at 37°C in 5% CO_2_, with protein transport inhibitor, GolgiStop (BD) added two hours after initiation of incubation. Stimulated splenocytes were then stained for lymphocyte surface markers CD8β and CD4, fixed with CytoFix (BD), permeabilized, and stained for intracellular accumulation of IFN-γ and TNF-α. Fluorescent-conjugated antibodies against mouse CD8β antibody (clone H35-17.2, ThermoFisher), CD4 (clone RM4-5, BD), IFN-γ (clone XMG1.2, BD), and TNF-α (clone MP6-XT22, BD) and staining was performed in the presence of unlabeled anti-CD16/CD32 antibody (clone 2.4G2). Flow cytometry was performed using a Beckman-Coulter Cytoflex S flow cytometer and analyzed using Flowjo Software.

### ELISpot assay

ELISpot assays were used to detect cytokines secreted by splenocytes from inoculated mice. Fresh splenocytes were used on the same day, as were cryopreserved splenocytes containing lymphocytes. The cells (2-4 × 10^5^ cells per well of a 96-well plate) were added to the ELISpot plate containing an immobilized primary antibodies to either IFN-γ or IL-4 (BD), and were exposed to various stimuli (e.g. control peptides, target peptide pools/proteins) comprising 2 μg/mL peptide pools or 10 μg/mL protein for 36-40 hours. After aspiration and washing to remove cells and media, extracellular cytokine was detected by a secondary antibody to cytokine conjugated to biotin (BD). A streptavidin/horseradish peroxidase conjugate was used detect the biotin-conjugated secondary antibody. The number of spots per well, or per 2-4 × 10^5^ cells, was counted using an ELISpot plate reader.

### ELISA for detection of antibodies

For antibody detection in sera from inoculated mice, ELISAs specific for spike and nucleocapsid antibodies, as well as for IgG subtype (IgG1, IgG2a, IgG2b, and IgG3) antibodies were used. A microtiter plate was coated overnight with 100 ng of either purified recombinant SARS-CoV-2 S-FTD (full-length S with fibritin trimerization domain, constructed and purified in-house by ImmunityBio), SARS-CoV-2 S RBD (Sino Biological, Beijing, China; Cat # 401591-V08B1-100) or purified recombinant SARS-CoV-2 nucleocapsid (N) protein (Sino Biological, Beijing, China; Cat # 40588-V08B) in 100 μL of coating buffer (0.05 M Carbonate Buffer, pH 9.6). The wells were washed three times with 250 μL PBS containing 1% Tween 20 (PBST) to remove unbound protein and the plate was blocked for 60 minutes at room temperature with 250 μL PBST. After blocking, the wells were washed with PBST, 100 μL of diluted serum samples were added to wells, and samples incubated for 60 minutes at room temperature. After incubation, the wells were washed with PBST and 100 μL of a 1/5000 dilution of anti-mouse IgG HRP (GE Health Care; Cat # NA9310V), or anti-mouse IgG_1_ HRP (Sigma; Cat # SAB3701171), or anti-mouse IgG_2a_ HRP (Sigma; Cat # SAB3701178), or anti-mouse IgG_2b_ HRP (Sigma; catalog# SAB3701185), or anti-mouse IgG_3_ HRP conjugated antibody (Sigma; Cat # SAB3701192) was added to wells. For positive controls, a 100 μL of a 1/5000 dilution of rabbit anti-N IgG Ab or 100 μL of a 1/25 dilution of mouse anti-S serum (from mice immunized with purified S antigen in adjuvant) were added to appropriate wells. After incubation at room temperature for 1 hour, the wells were washed with PBS-T and incubated with 200 μL o-phenylenediamine-dihydrochloride (OPD substrate (Thermo Scientific Cat # A34006) until appropriate color development. The color reaction was stopped with addition of 50 μL 10% phosphoric acid solution (Fisher Cat # A260-500) in water and the absorbance at 490 nm was determined using a microplate reader (SoftMax® Pro, Molecular Devices).

### Calculation of relative μg amounts of antibodies

A standard curve of IgG was generated and absorbance values were converted into mass equivalents for both anti-S and anti-N antibodies. Using these values, we were able to calculate that hAd5 S-Fusion + N-ETSD vaccination generated a geometric mean value of 5.8 μg S-specific IgG and 42 μg N-specific IgG per milliliter of serum.

### cPass™ Neutralizing Antibody Detection

The GenScript cPass™ (https://www.genscript.com/cpass-sars-cov-2-neutralization-antibody-detection-Kit.html) for detection of neutralizing antibodies was used according to the manufacturer’s instructions.^44^ The kit detects circulating neutralizing antibodies against SARS-CoV-2 that block the interaction between the S RBD with the ACE2 cell surface receptor. It is suitable for all antibody isotypes and appropriate for use with in animal models without modification.

### Vero E6 cell neutralization assay

All aspects of the assay utilizing virus were performed in a BSL3 containment facility according to the ISMMS Conventional Biocontainment Facility SOPs for SARS-CoV-2 cell culture studies. Vero e6 kidney epithelial cells from *Cercopithecus aethiops* (ATCC CRL-1586) were plated at 20,000 cells/well in a 96-well format and 24 hours later, cells were incubated with antibodies or heat inactivated sera previously serially diluted in 3-fold steps in DMEM containing 2% FBS, 1% NEAAs, and 1% Pen-Strep; the diluted samples were mixed 1:1 with SARS-CoV-2 in DMEM containing 2% FBS, 1% NEAAs, and 1% Pen-Strep at 10,000 TCID50/mL for 1 hr. at 37°C, 5% CO^2^. This incubation did not include cells to allow for neutralizing activity to occur prior to infection. The samples for testing included sera from the four mice that showed > 20% inhibition of ACE2 binding in cPass, pooled sera from those four mice, sera from a COVID-19 convalescent patient, and media only. For detection of neutralization, 120 μL of the virus/sample mixture was transferred to the Vero E6 cells and incubated for 48 hours before fixation with 4% PFA. Each well received 60 μL of virus or an infectious dose of 600 TCID50. Control wells including 6 wells on each plate for no virus and virus-only controls were used. The percent neutralization was calculated as 100-((sample of interest-[average of “no virus”])/[average of “virus only”])*100) with a stain for CoV-2 Np imaged on a Celigo Imaging Cytometer (Nexcelom Bioscience).

